# Jackknife model averaging prediction methods for complex phenotypes with gene expression levels by integrating external pathway information

**DOI:** 10.1101/447706

**Authors:** Xinghao Yu, Lishun Xiao, Ping Zeng, Shuiping Huang

## Abstract

**Motivation:** In the past few years many novel prediction approaches have been proposed and widely employed in high dimensional genetic data for disease risk evaluation. However, those approaches typically ignore in model fitting the important group structures or functional classifications that naturally exists in genetic data.

**Methods:** In the present study, we applied a novel model averaging approach, called Jackknife Model Averaging Prediction (JMAP), for high dimensional genetic risk prediction while incorporating KEGG pathway information into the model specification. JMAP selects the optimal weights across candidate models by minimizing a cross-validation criterion in a jackknife way. Compared with previous approaches, one of the primary features of JMAP is to allow model weights to vary from 0 to 1 but without the limitation that the summation of weights is equal to one. We evaluated the performance of JMAP using extensive simulation studies and compared it with existing methods. We finally applied JMAP to five real cancer datasets that are publicly available from TCGA.

**Results:** The simulations showed that, compared with other existing approaches, JMAP performed best or are among the best methods across a range of scenarios. For example, among 14 out of 16 simulation settings with PVE=0.3, JMAP has an average of 0.075 higher prediction accuracy compared with gsslasso. We further found that in the simulation the model weights for the true candidate models have much smaller chances to be zero compared with those for the null candidate models and are substantially greater in magnitude. In the real data application, JMAP also behaves comparably or better compared with the other methods for both continuous and binary phenotypes. For example, for the COAD, CRC and PAAD data sets, the average gains of predictive accuracy of JMAP are 0.019, 0.064 and 0.052 compared with gsslasso.

**Conclusion:** The proposed method JMAP is a novel method that can provide more accurate phenotypic prediction while incorporating external useful group information.

## Introduction

Due to the rapid development of biotechnology [1–4], a large number of high-throughput and low-cost gene sequencing data have been generated and provide a broad space to investigate the association between genetic factors and complex diseases/disorders[5–14]. The great succeed of association studies further promotes the genetic risk prediction and evaluation for complex phenotypes by incorporating into omics information [15–18]. Due to the high dimensional problem that the number of genetic markers is much larger than the sample size, one of the greatest challenges for genetic risk prediction is that it is difficult to apply traditional statistical methods for classification and prediction in large scale molecular omics data. In the past few years, developing prediction methods that can efficiently model high dimensional genetic data has been an active area and attached much research attention; and a series of novel prediction approaches have been proposed and widely employed for disease risk evaluation or gene expression imputation [19–25]. However, those approaches typically ignore in model fitting the important group structures or functional classifications that naturally exist in genetic data. For example, it is well known that single nucleotide polymorphisms (SNPs) can be divided into groups in terms of functional annotations or genes, and genes in turn can be grouped into pathways due to the shared biological function. It has been shown that incorporating such useful group/functional information into model fitting can substantially boost statistical power in genetic association studies and can facilitate our understanding of the genetic architecture of disease variation by heritability partition [25–33]. One widely-used group source is the pathway information in the Kyoto Encyclopedia of Genes and Genomes (KEGG) [34, 35], which integrates information on genomic, chemical and system functions and groups genes with highly related sequences by analyzing the sequence similarity of genes.

Besides in genetic association studies and heritability estimation, it has been also shown that the prediction accuracy can be improved by leveraging grouped functional information in genetic risk evaluation with large scale omics data [36–38]. For example, Tang et al [38] recently designed a group spike-and-slab Lasso generalized linear model (gsslasso) that combined KEGG pathway information into model fitting and demonstrated that, compared with Lasso [39], the average gains of prediction accuracy [measured by area under the curve (AUC)] of gsslasso were about 4.5% for sarcoma, 4.6% for ovarian cancer and about 1.6% for breast cancer by leveraging gene expression data available from The Cancer Genome Atlas (TCGA) [40]. Motivated by those results, in this study we employ a novel model averaging approach [41, 42] for genetic risk prediction while incorporating KEGG pathway information into the model specification. The proposed model averaging approach selects the optimal weights across candidate models by minimizing a cross-validation criterion in a jackknife way. We thus refer to the present method as **J**ackknife **M**odel **A**veraging **P**rediction (JMAP). We use extensive simulation studies to evaluate the performance of JMAP and compare it with existing methods. Finally, we apply JMAP to five real cancer datasets that are publicly available from TCGA. To construct candidate prediction models, we divide genes in terms of the KEGG pathway information [34, 35].

## Methods and Materials

### Overview of the JMAP Method

We first present an overview of JMAP here; the detailed description of JMAP is shown in the Supplementary Text. Briefly, JMAP consists of two-step model fitting procedures: **(i)** in the first step, we group the predictors (e.g. genome-wide gene expression levels) and build a series of candidate linear prediction models with the gene expression measurements available for various groups; **(ii)** in the second step, we look for a suitable model weight vector for averaging across the candidate models to perform a pooled model prediction. One of the primary features of JMAP is to allow model weights to vary from 0 to 1 but without the limitation that the summation of weights is necessarily equal to one [41, 42]. As we will see, this weight relaxation is important and critical, resulting in an effective improvement of the prediction accuracy. JMAP has been implemented within an R function freely available at https://github.com/biostatpzeng.

### Simulations and real data applications

#### Simulation settings

We next carried out extensive simulations to evaluate the prediction performance of JMAP. To make the simulation settings as real as possible, we used gene expression levels obtained from an existing TCGA data set of breast cancer (see below for further information about this data). For simplicity, we extracted the expression levels for 6,000 randomly selected genes and 500 breast cancer patients and simulated phenotypes using the following model

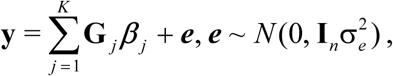

where *K* is the total number of groups (or pathways); **G***_j_* is an *n*×*m_j_* genetic matrix for *m_j_* genes in group *j* with *n* the sample size, ***β****_j_* is an *m_j_*-dimensional vector of effects sizes (here *n*=500); **I***_n_* is an *n*×*n* identity matrix; and ***e*** is an *n*-dimensional vector of independently and normally distributed residuals with variance 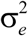. We considered four scenarios with different group partitions. In scenarios 1-3, genes were sequentially divided into 50, 200 or 300 groups with approximately equal genes per each group; no overlapping of genes existed among groups. In scenario 4, we classified genes into 328 groups in terms of the KEGG pathway information (see below for details); note that, under this case the number of genes included in each group was not equal and ∼21% genes belonged to multiple pathways. Then, following [38], in each scenario we randomly selected five out of all *K* groups (*K*=50, 200, 300 or 328 as defined above) and generated: (**I**) the effect sizes ***β****_l_* (*l*=1, 2, 3, 4 and 5) in each of the selected groups followed a normal distribution with mean zero and the same variance (say 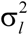). Under this case, all the genes in the five groups had non-zero effect sizes; **(II)** unlike the case **I**, here we assumed that only the genes in the first two groups had non-zero effect sizes and half of the genes in the last three group had non-zero effect sizes; **(III)** instead of assuming equal proportion of non-zero effect sizes in the last three groups, we set the proportion of non-zero effect sizes to be 80%, 50% and 20%, respectively; **(IV)** in this case, we set the proportion of non-zero effect sizes to be 90%, 70%, 50%, 30% and 10% for the five groups, respectively. The variance parameters 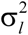 and 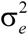 were carefully chosen to ensure that **y** had unit variance asymptotically and the phenotypic variance explained (PVE) by genetic component was 0.3, 0.5 or 0.8 in each case, respectively. The effect sizes for the unselected gene groups were set to zero.

#### Real data applications

We now applied JMAP to five cancer data sets publicly available from TCGA [40], including the breast cancer (BRCA), the colon and rectal cancer (CRC), the colon cancer (COAD), the lower grade glioma (LGG) and the pancreatic cancer (PAAD). We downloaded both the clinical data and RNAseq gene expression levels for those cancers from UCSC Xena (https://xenabrowser.net/). For each cancer, we first merged the clinical data and gene expression levels measured from primary cancer tissue; then we removed genes with more than 50% zero expressions and standardized the remaining gene expression levels. The used data sets in this study were summarized in Table 1. Following previous studies [12, 43, 44], for the five cancers we first used the age at initial pathologic diagnosis (i.e. onset age) as phenotypes because the age of onset is an important indicator that the cancer is likely more commonly genetic in origin. Besides the age of onset, we also applied our method to the count of positive lymph nodes identified through hematoxylin and eosin staining light microscopy for BRCA (LYM of BRCA). The number of lymph node metastases and whether there is lymphatic metastasis have a strong adverse effect on the prognosis of tumor patients [45]. For this variable, we first regressed out the influence of pathologic stage and radiation therapy using a Poisson model, and kept the resulting residual as the phenotype. Both the age of onset and the residual of LYM were continuous and were quantile-normalized to a standard normal distribution before prediction analysis. Additionally, we also considered two binary phenotypes (assumed to be labeled as 0 and 1): the occurrence of new tumor event after initial treatment for LGG (Recurrence of LGG) and the lymphatic metastasis for BRCA (Metastasis of BRCA). Rather than fitting logistic candidate regressions for the binary phenotypes [46], we instead treated them as continuous outcomes and directly fitted linear candidate models. We will further discuss this issue later.

**Table 1.**
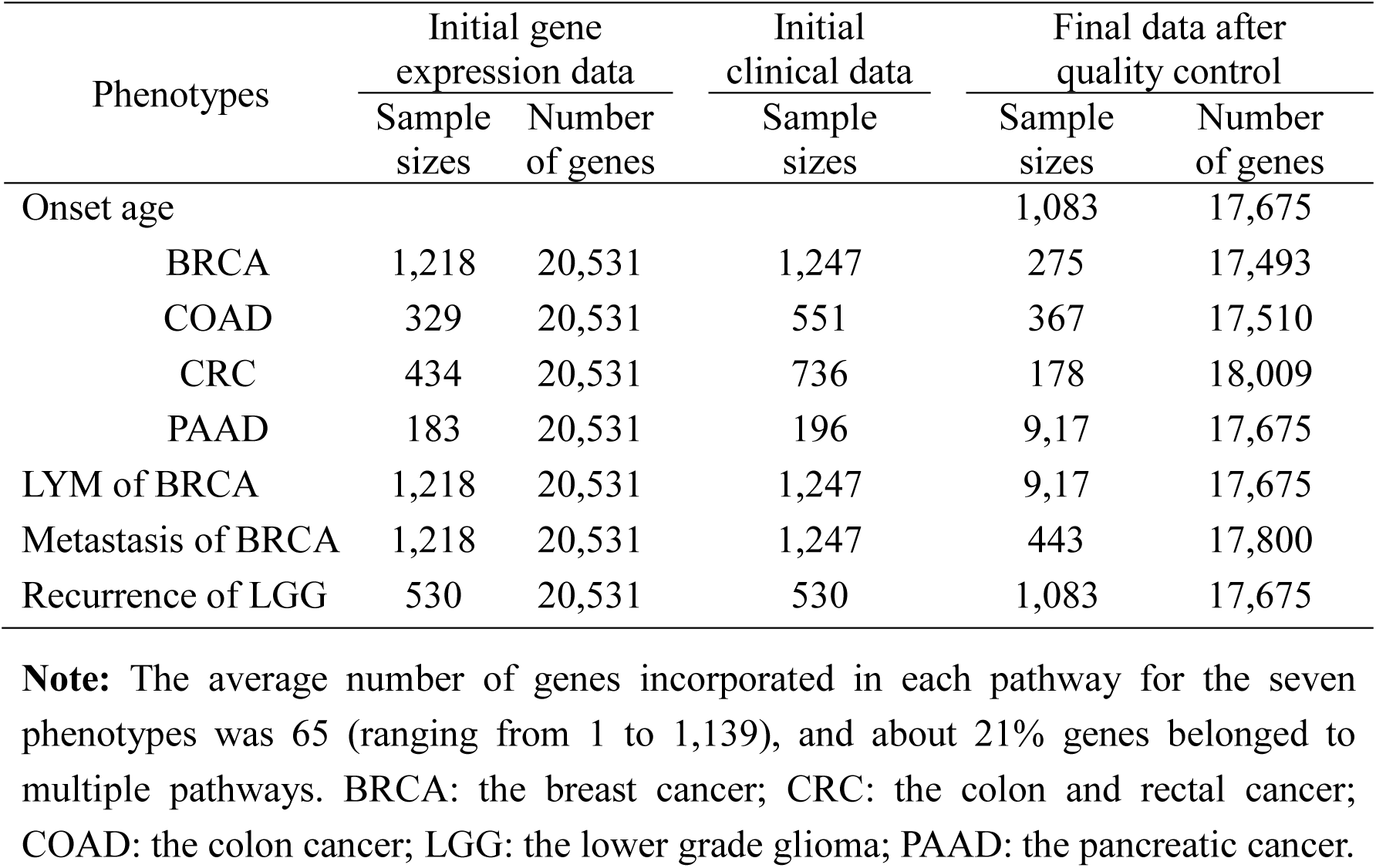
Sample sizes and the number of genes for each cancer in the TCGA data set used in our analysis

| Phenotypes | Initial gene expression data |  | Initial clinical data | Final data after quality control |  |
| --- | --- | --- | --- | --- | --- |
|  | Sample sizes | Number of genes | Sample sizes | Sample sizes | Number of genes |
| Onset age |  |  |  | 1,083 | 17,675 |
| BRCA | 1,218 | 20,531 | 1,247 | 275 | 17,493 |
| COAD | 329 | 20,531 | 551 | 367 | 17,510 |
| CRC | 434 | 20,531 | 736 | 178 | 18,009 |
| PAAD | 183 | 20,531 | 196 | 9,17 | 17,675 |
| LYM of BRCA | 1,218 | 20,531 | 1,247 | 9,17 | 17,675 |
| Metastasis of BRCA | 1,218 | 20,531 | 1,247 | 443 | 17,800 |
| Recurrence of LGG | 530 | 20,531 | 530 | 1,083 | 17,675 |

**Note:** The average number of genes incorporated in each pathway for the seven phenotypes was 65 (ranging from 1 to 1,139), and about 21% genes belonged to multiple pathways. BRCA: the breast cancer; CRC: the colon and rectal cancer; COAD: the colon cancer; LGG: the lower grade glioma; PAAD: the pancreatic cancer.

### Model comparison and implementation

For the simulated data the genes were divided into 50, 200, 300 or 328 groups under various scenarios as mentioned before. For the real data sets, we mapped the genes to KEGG pathways by R package clusterProfiler (version 3.8.1) after matching gene symbols to Entriz ids [47], and divided the genes into 328 pathway groups. For both simulated and real data sets, following [22] we performed 100 Monte Carlo cross validation (MCCV) data splits by randomly selecting 80% samples as training data and the remaining 20% as test data. We fitted the prediction models in the training data and evaluated the performance in the test data with correlation coefficient (R) for the continuous phenotypes (i.e. onset age of BRCA, COAD, CRC, LGG and PAAD, and LYM of BRCA) or AUC for the binary phenotypes (i.e. Tumor of LGG and Metastasis of BRCA).

The competing methods included Lasso [39], Elastic Net (ENET) [48] and gsslasso [38]. For both Lasso and ENET, we implemented them via the R package glmnet (version 2.0-16), selected the optimal penalty parameters in Lasso and ENET using 100-fold cross validation, and set α=0.50 in ENET as done in [49]. For gsslasso, we implemented it via the R package BhGLM (version 1.1.0). Following [38] we selected the optimal penalty parameter of gsslasso by setting the slab scale (denoted by *s*_1_) to 1, calculated the accuracy of prediction for a series values for the spike scale (denoted by *s*_0_) (i.e. *s*_0_=0.01×*m*, *m*=0.1, 1, 2, …, 9) and chose the optimal value for *s*_0_ that resulted in a highest prediction. We solved the quadratic problem in JMAP as shown in Equations (7)-(8) (see Supplementary Text) by using the optim function in R software. We further contrasted the prediction performance of all other methods with that of JMAP by taking the difference of *R* or AUC between the other methods and JMAP. Therefore, an *R* or AUC difference below zero suggests worse performance than JMAP.

## Results

### Results of the simulation studies

The simulation results for the difference of *R* with PVE=0.3 are shown in Figure 1 with the original *R* values shown in Figure S1. There are 16 combinations presented in Figure 1. Compared with other existing approaches (i.e. Lasso, ENET and gsslasso), we find that, except two situations, JMAP performed best or are among the best methods in most of the combinations (14 out of 16). For example, among those 14 settings, JMAP has an average of 0.075 higher prediction accuracy compared with gsslasso, with the difference of *R* ranging from 0.023 to 0.116. In the setting with 200 groups in scenario **I** (where all the genes in the five groups had non-zero effect sizes), JMAP behaves slightly worse than Lasso (0.012 lower) and ENET (0.013 lower), but is better than gsslasso (0.056 higher). In the setting with 300 group in scenario **III** (where the genes among the first two groups had non-zero effect sizes, but some of the genes in the rest three groups are null with various null proportions), all the three competitive methods (i.e. Lasso, ENET and gsslasso) have a higher prediction accuracy relative to JMAP. The simulation results for PVE=0.5 and 0.8 are displayed in Figure S2-S5 in the Supplementary Text; we observe the similar pattern that JMAP performs better or is as good as other competing methods in most of the simulated settings. We further check the estimated weights for the candidate models in all the scenarios and find that the weights for the true candidate models (i.e. those with non-zero effect sizes) have much smaller chances to be zero compared with those for the null candidate models and are substantially greater in magnitude (Table S1).

**Figure 1.**
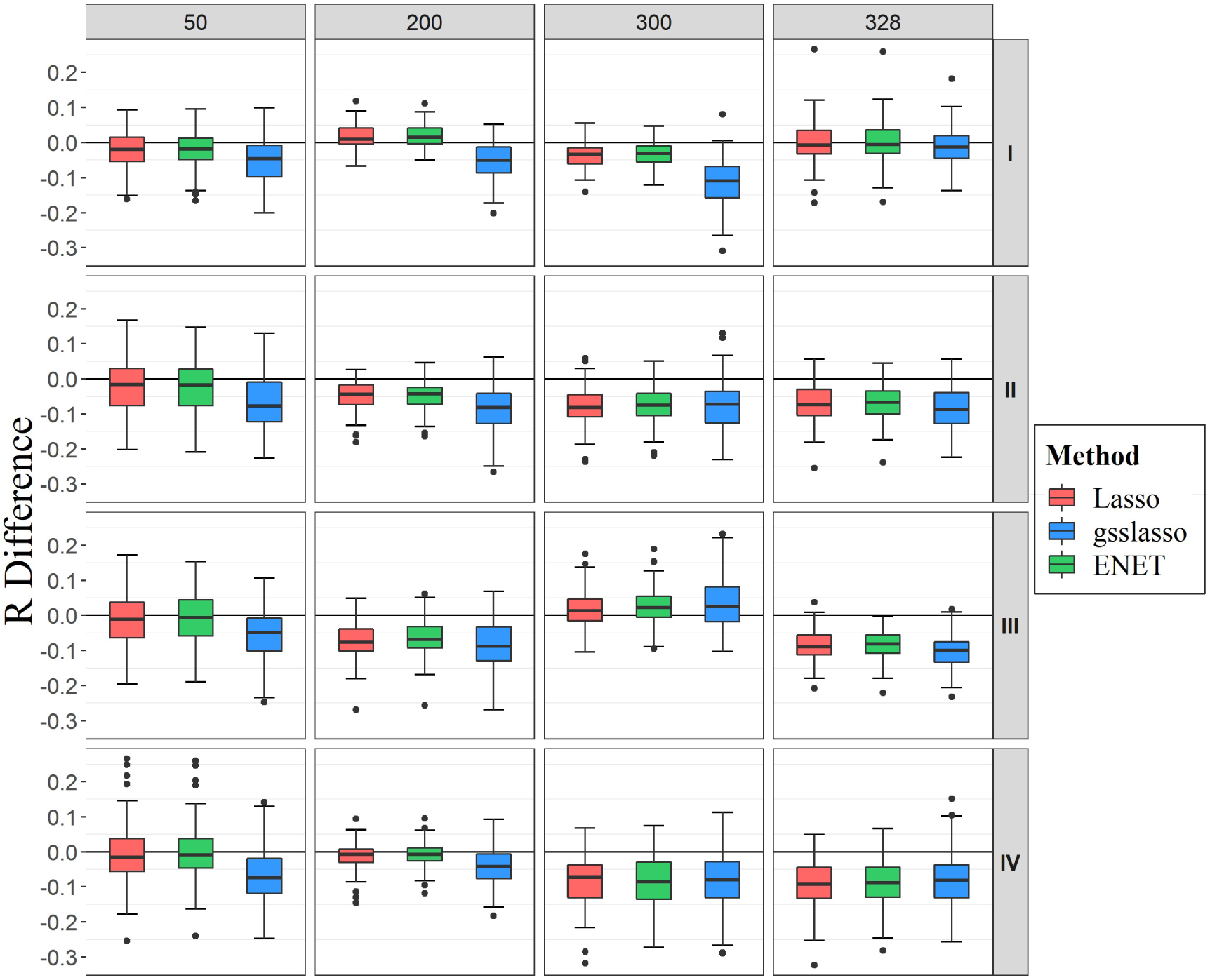
Comparison of predictive performance of three models with JMAP with PVE=0.3. Performance is measured by *R* difference with respect to JMAP; therefore, a negative value (i.e. values below the horizontal line) indicates worse performance than JMAP. In each setting, five groups with non-zero effect sizes were selected; I represents the settings where all the genes in the five groups had non-zero effect sizes; II represents the settings where only the genes in the first two groups had non-zero effect sizes and half of the genes in the last three group had non-zero effect sizes; III represents the settings where the effect sizes of the first two groups were non-zero and the proportion of non-zero effect sizes in the last three groups was 80%, 50% or 20%; IV represents the settings where the proportion of non-zero effect sizes in the five groups was 90%, 70%, 50%, 30% or 10%. The predictive performance was assessed across 100 replicates in each scenario.

### Results of the real data applications

Now, we turn to the real application of the TCGA data (Table 1). The results of *R* difference for continuous phenotypes and AUC difference for binary phenotypes are presented in Figure 2. Figure 2A shows the predictive performance of other three methods compared with JMAP for five continuous phenotypes. It can be seen that totally JMAP performs comparably or better compared with the other methods. For example, for the age onset in the COAD, CRC and PAAD data sets, JMAP has the highest predictive power, followed by gsslasso. Compared with gsslasso, in these three data sets the gains of predictive accuracy of JMAP are 0.019, 0.064 and 0.052, respectively. However, Lasso, gsslasso and ENET have higher prediction accuracy for the initial age of onset of breast cancer patients. For LYM of BRCA, the four methods behave very consistently, while JMAP has a slightly small advantage over the rest ones. With regard to the two binary phenotypes (Figure 2B), JMAP still maintains stable and robust predictive performance. For metastasis of BRCA, gsslasso performs the best, following by Lasso and ENET. For recurrence of LGG; JMAP performs best and has a 0.087 higher predictive accuracy compared with gsslasso.

**Figure 2.**
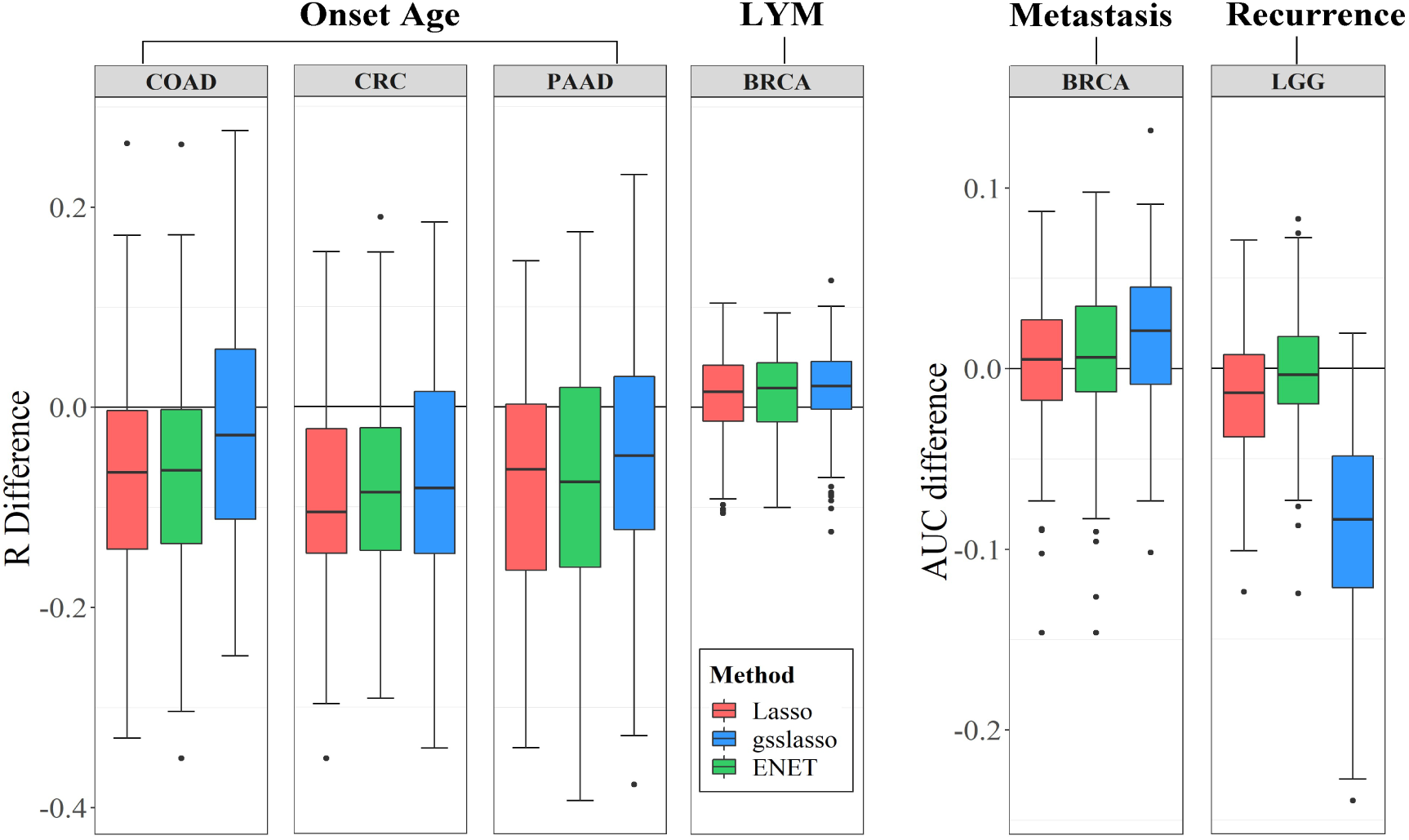
Comparison of predictive performance of three models with JMAP for seven phenotypes from the TCGA data sets. Performance is measured by *R* (or AUC) difference with respect to JMAP; therefore, a negative value (i.e. values below the horizontal line) indicates worse performance than JMAP. The predictive performance was assessed across 100 MCCV replicates. BRCA: the breast cancer; CRC: the colon and rectal cancer; COAD: the colon cancer; LGG: the lower grade glioma; PAAD: the pancreatic cancer.

## Discussion

In the present study we have employed a novel statistical method, JMAP, for genetic prediction and evaluation of complex phenotypes from the publicly available TCGA data sets. Traditionally, the classical model averaging methods first build a series of candidate models with various degrees of model complexity; then combine all the candidate models together to boost the prediction performance by specifying greater weights onto better models; and require the summation of the model weights is equal to one [41, 50, 51]. However, unlike those previous methods, MAP relaxes the constraint of summing the weights of candidate models up to one. By removing this restriction and including genetic pathway information, as we have demonstrated in the simulations and real data applications, JMAP has shown higher prediction accuracy compared existing approaches. Furthermore, it is natural to examine whether the weight restriction can be further relaxed to allow them to vary between −1 and 1 [46]. However, we found that this further relaxing may be not beneficial for improving the prediction performance, leading to low accuracy of genetic prediction (Figure S7). Additionally, because each candidate model is fitted with ordinary lease squares method which leads to an analytical solution for the effect sizes, and because the weight estimation is optimized through a constrained quadratic manner, JMAP is thus computationally efficient and can be easily scalable to the high dimensional genetic risk prediction problem. For example, in our real data applications, it takes only about 18, 21 and 200 seconds on average for the COAD, PAAD and BRCA data sets, respectively.

In practice, the candidate models for model averaging are typically established in terms of prior knowledge or expert viewpoints and the number of the candidate models (i.e. *K* in our study) is assumed to be uncertain. To address this problem, Ando and Li [42] recently proposed first to partition predictors (equivalent to genes in our study) based on the marginal correlation magnitude between each predictor and the response; and then adaptively prepared for candidate model for each partition. This strategy is a flexible way and avoids the requirement of external information; while it may be suboptimal if there is informative prior information that can be utilized. In contrast, in our study we explicitly preassigned the number of candidate models for JMAP. Indeed, using simulations we have discovered that JMAP possessed consistently good prediction performance across various candidate model partitions (Figure S8). In our real data applications, we also directly built the candidate models for JMAP based on useful KEGG pathway information which characterizes the biological functions for various sets of genes [34, 35] and can result in each candidate model having unique strength in capturing certain aspects of prediction ability. Applying external informative pathways to establish candidate models in JAMP can lead to at least three benefits: **(i)** it does not need to search for the appropriate number of candidate models by partitioning all the genes; thus, it is computationally faster; **(ii)** because relying on previously well-validated pathway information, the established candidate models is more biologically meaningful; **(iii)** finally, the marginal correlation way typically groups a given gene into only one candidate model [42]; while in practice a gene often can be involved in multiple pathways and will be thus included into several candidate models; e.g. in our analysis about 21% genes can be grouped into at least two pathways. More generally, under the context of model averaging JAMP can naturally handle the overlapping group structures — a phenomenon that is frequently encountered in pathway-based data analyses [52]. It has been shown that efficiently incorporating the overlapping group structures into model fitting can raise the prediction performance [38]. Hence, JAMP has the potential for further enhancing prediction accuracy.

It is worth noting that in the candidate model of JMAP the least squares estimate in Equation (2) (Supplementary Text) is ill-conditional when the number of genetic markers is larger than the sample size for some genes. For example, in our analysis there are 5.5% and 5.2% pathways with the number of genes greater than the sample sizes for the PAAD and COAD data sets, respectively. Under this situation, by borrowing the idea of ridge regression [53, 54], we have attempted to add a non-negative constant δ into the estimates; i.e. replacing 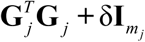 with 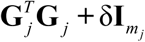 (see Equation 2 in the Supplementary Text). In the present study we primarily set 5 to be one and found that JMAP is robust against with regard to various values of 5 with simulations (Figure S9). We emphasize that this is an ad hoc modification which has no clear theoretical foundation. Further investigation of JMAP under the context that the dimension of candidate model is larger than the sample size is an important and interesting topic and is our next research direction.

Finally, the current version of JMAP described in our study is constructed only for continuous phenotypes. Extending model averaging from linear to nonlinear regression under the high dimensional situations was recently investigated [46]. However, although not mentioned, an explicit model assumption in their study is that the number of the predictors in each candidate generalized linear model should be much less than the sample size to ensure the estimates can be identifiable. Therefore, their methods cannot be applied to our case where the number of the genes for some candidate models is easy to be greater than the sample size as mentioned before. Thus, in our real data application we had to directly fit linear candidate models for binary phenotypes by treating them as continuous values following previous studies [19–21, 23]. Theoretically, modeling binary data with linear models can be justified by the fact that the linear model can be viewed as a first order Taylor approximation to the generalized linear model; and this approximation is accurate when the effect size is weak and small [19] — a condition which generally satisfies because it has been shown that most complex phenotypes are polygenic and are influenced by many genetic variants with small effect sizes [55]. Nevertheless, extending the JMAP model for application to non-continuous phenotypes in high dimensional prediction problems warrants more explorations.

## Abbreviations

JMAP: Jackknife model averaging prediction
ENET: Elastic net
gsslasso: Group spike-and-slab lasso
AUC: Area under the curve
KEGG: Kyoto Encyclopedia of Genes and Genomes

## Acknowledgements

We acknowledge the contributions of TCGA Research Network.

## Availability of data and materials

The TCGA data is publicly available from https://xenabrowser.net/. The BhGLM software is available from http://github.com/nyiuab/BhGLM, the glmnet package is available from https://cran.r-project.org/web/packages/glmnet/index.html.

## Authors’ contributions

PZ and SH conceived and designed the experiment. XY and PZ cleared up, analyzed and interpreted the data sets, XY, LX and PZ wrote the manuscript. All authors read and approved the final manuscript.

## Consent for publication

Not applicable.

## Ethics approval and consent to participate

Not applicable.

